# Dihydrothiazolo ring-fused 2-pyridone antimicrobial compounds treat *Streptococcus pyogenes* skin and soft tissue infection

**DOI:** 10.1101/2024.01.02.573960

**Authors:** Zongsen Zou, Chloe L. P. Obernuefemann, Pardeep Singh, Jerome S. Pinkner, Wei Xu, Taylor M. Nye, Karen W. Dodson, Fredrik Almqvist, Scott J. Hultgren, Michael G. Caparon

## Abstract

We have developed GmPcides from a peptidomimetic dihydrothiazolo ring-fused 2-pyridone scaffold that have antimicrobial activities against a broad-spectrum of Gram-positive pathogens. Here we examine the treatment efficacy of GmPcides using skin and soft tissue infection (SSTI) and biofilm formation models by *Streptococcus pyogenes*. Screening our compound library for minimal inhibitory (MIC) and minimal bactericidal (MBC) concentrations identified GmPcide PS757 as highly active against *S. pyogenes*. Treatment of *S. pyogenes* biofilm with PS757 revealed robust efficacy against all phases of biofilm formation by preventing initial biofilm development, ceasing biofilm maturation and eradicating mature biofilm. In a murine model of *S. pyogenes* SSTI, subcutaneous delivery of PS757 resulted in reduced levels of tissue damage, decreased bacterial burdens and accelerated rates of wound-healing, which were associated with down-regulation of key virulence factors, including M protein and the SpeB cysteine protease. These data demonstrate that GmPcides show considerable promise for treating *S. pyogenes* infections.

## INTRODUCTION

The emergence of antibiotic resistance threatens healthcare and agriculture systems worldwide and raises the prospect of a post-antibiotic era. Several factors, including the overuse and misuse of antibiotics, and exposure to environmental reservoirs of antibiotic-resistant bacteria have contributed to rising rates of antibiotic resistance (*1, 2*). A slowdown in new antimicrobial drug development has resulted in reliance on existing antimicrobials, which when combined with poor antibiotic stewardship, has further exacerbated the development of resistance (*3, 4*). Thus, there is an urgent need to develop new antibiotics that are recalcitrant to resistance development to combat multidrug-resistant pathogens.

Towards this goal, we have developed GmPcides, a new class of synthetic compounds based on a peptidomimetic dihydrothiazolo ring-fused 2-pyridone scaffold. Rational alteration at positions C-2, C-7 and C-8 of the central fragment via various synthetic methodologies has resulted in the development of GmPcides with enhanced antibacterial and drug-like activities (*5*). Among these is GmPcide PS757 with robust bacteriostatic and bactericidal activities against a broad range of multidrug-resistant Gram-positive pathogens (*5*), including vancomycin-resistant *Enterococcus faecalis* (VRE), methicillin-resistant *Staphylococcus aureus* (MRSA), multidrug-resistant *Streptococcus pneumoniae*, clindamycin-resistant *Streptococcus agalactiae* and erythromycin-resistant *Streptococcus pyogenes*, all of which are classified as serious or concerning threats by the Centers for Disease Control and Prevention (CDC) (*6*). Given the broad-spectrum antibacterial activities of GmPcides against Gram-positive pathogens, the next phase of development is to assess their efficacy for treatment using well-characterized models of Gram-positive infection and biofilm formation.

For this study, we have focused on *S. pyogenes*, a Gram-positive pathogen responsible for over 500,000 deaths per year, a global burden that approaches that of rotavirus and measles (*7*). In humans, *S. pyogenes* can cause a wide range of diseases, ranging from mild to severe, including pharyngitis (strep throat), scarlet fever, toxic shock syndrome, acute glomerulonephritis and rheumatic fever (*8–11*). Of particular concern are skin and soft tissue infections (SSTI) that range from superficial infection of the epidermis such as impetigo, to severe invasive infections of the dermis and deeper tissues including cellulitis and necrotizing fasciitis (the “flesh-eating disease”) (*12, 13*). Despite its sensitivity to many different antibiotics, including β-lactams, treatment of *S. pyogenes* invasive infection is complicated by factors that include its ability to form biofilm and its ability to secrete a myriad number of toxins. Biofilm is an adherent and structured community of bacteria growing within an extracellular polymeric substance (EPC) that enhances the community’s ability to resist killing by antibiotics (*14–16*). Toxins produced by *S. pyogenes* play a major role in damaging host tissue and include several membrane-disruptive hemolysins, immuno-modulating superantigens, plasminogen activators, host cell adhesins, complement-modulating proteins, specific and non-specific proteases and multiple other degradative enzymes. Since tissue damage impacts the efficiency of antibiotics to kill *S. pyogenes*, treatment strategies for necrotizing disease often include an antibiotic that inhibits expression of tissue-damaging toxins in combination with an antibiotic that targets bacterial growth. The latter typically includes a β-lactam (e.g., piperacillin/tazobactam), while the former includes clindamycin (*13, 17, 18*), which at sublethal concentrations *in vitro,* inhibits expression of several tissue-damaging toxins. However, the use of clindamycin is now threatened by increasing rates of resistance in healthcare settings (*17–19*).

In the present study, we extensively characterized the therapeutic properties of GmPcide PS757 using well-characterized biofilm and murine SSTI models of *S. pyogenes* infection. Our results demonstrate that PS757 was effective against all stages of bacterial growth and biofilm formation *in vitro*, and improved treatment outcomes in a murine SSTI model by decreasing bacterial burdens, reducing levels of tissue damage, attenuating inflammation and accelerating the rate of wound-healing. Sublethal exposure to GmPcides reduced virulence factor expression, including expression of two major virulence factors, the surface associated M protein and the secreted SpeB cysteine protease. Together, these results suggest that GmPcides have considerable promise for preventing and treating *S. pyogenes* infections.

## RESULTS

### GmPcide PS757 Induces Nucleoid and Cell Wall Abnormalities

In our previous study, we have identified a GmPcide PS757 with robust antimicrobial activity against a wide range of multidrug-resistant Gram-positive pathogens, including the *S. pyogenes* that was investigated in the present study (*5*). The minimum inhibitory concentration (MIC) of 0.78 µM PS757 against *S. pyogenes* HSC5 strain as determined in the previous study (*5*) demonstrates its robust bacteriostatic activity. In the present work, we further determined its minimum bactericidal concentration (MBC) against streptococcal cells as 1.56 µM, revealing its effective bactericidal activity as well (Table 1). Moreover, exponential and stationary phases *S. pyogenes* HSC5 cells treated under a bactericidal concentration (20 µM) of PS757 for 12 hours were observed with > 6 log and > 5 log CFU reductions, respectively (Table 1 and fig. S1), demonstrating its antimicrobial efficacy against both non-dividing and dividing streptococcal cells. A sublethal concentration of PS757 was also determined based on both growth curves (OD_600_) and growth yields (CFU) as 0.4 µM (fig. S2). Examination of PS757-treated cells by transmission electron microscopy (TEM) revealed that when compared to vehicle-treated (DMSO) cells, challenge with a sublethal concentration (0.4 µM) and with a bactericidal concentration (20 µM) both induced extensive nucleoid abnormalities characterized by a condensed and filamentous nucleoid structure (Fig. 1). In addition, cells treated at the bactericidal concentration (20 µM) displayed numerous small dense globular structures formed at the periphery of the cell wall (Fig. 1), suggesting that PS757 induces cell envelope abnormalities.

**Fig. 1.**
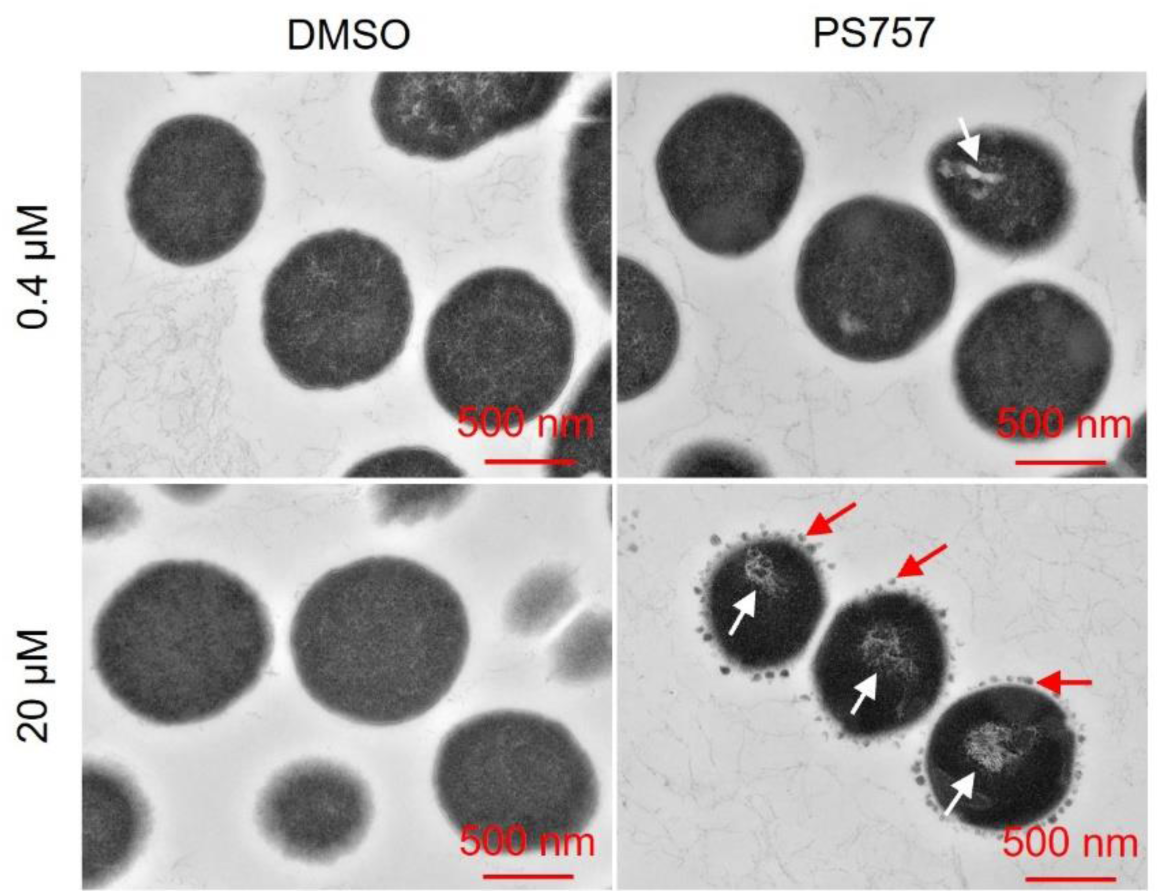
GmPcide PS757 treatment caused nucleoid and cell wall abnormalities in *S. pyogenes* cells. *S. pyogenes* HSC5 cells treated under both sublethal (0.4 µM) and bactericidal (20 µM) concentrations of PS757 exhibited nucleoid abnormality with altered nucleoid structure that was less dense and very filamentous (white arrows). *S. pyogenes* HSC5 cells treated under bactericidal (20 µM) concentration of PS757 were observed with small dense globular structures at periphery of bacterial cell wall (red arrows), suggesting that PS757 produced cell wall abnormalities.

**Table 1.** Antimicrobial activity (µM) of GmPcide PS757 against *S. pyogenes* HSC5.

| Strain | PS757 ($\mu\text{M}$) | | PS757 (20 $\mu\text{M}$) | |
| --- | --- | --- | --- | --- |
|  | MIC | MBC | Exponential | Stationary |
| *S. pyogenes* HSC5 | 0.78<sup>a</sup> | 1.56 | > 6 logCFU reduction | > 5 logCFU reduction |

### GmPcide PS757 Prevents *S. pyogenes* Biofilm Formation

We used an established model for *S. pyogenes* biofilm formation with BHI medium in a 96-well plate format to analyze the efficacy of GmPcides against biofilm. Planktonic growth was measured by OD_600_ (Fig. 2A) and biofilm formation determined using a standard crystal violet-staining assay (Fig. 2B). In the absence of GmPcide treatment, detectable biofilm began to accumulate at 4 hrs post-inoculation and continued to develop until reaching maturity at approx. 12 hrs (Fig. 2B). Cultures were then treated at 4, 7 and 24 hrs post-inoculation (fig. S3, A, B, and CA-C) with a series of different concentrations of PS757 to examine its activity against different phases of biofilm formation. During the initiation phase (4 hrs), PS757 at 0.7 µM and 1.0 µM was able to prevent both planktonic growth and biofilm formation (Fig. 2C). When added to developing biofilm at 7 hrs, 2.0 µM PS757 prevented further planktonic growth and at 5.0 µM arrested further maturation of biofilm (Fig. 2D). Finally, when used to treat a fully mature biofilm at 24 hrs, a concentration of 20 µM produced >90% decrease in cell viability when measured using a fluorescent vital stain and confocal microscopy after 5 hrs of treatment (Fig. 2E). In summary, PS757 was efficacious against all phases of *S. pyogenes* biofilm development including initiation and maturation, and had the ability to kill cells in a fully mature biofilm.

**Fig. 2.**
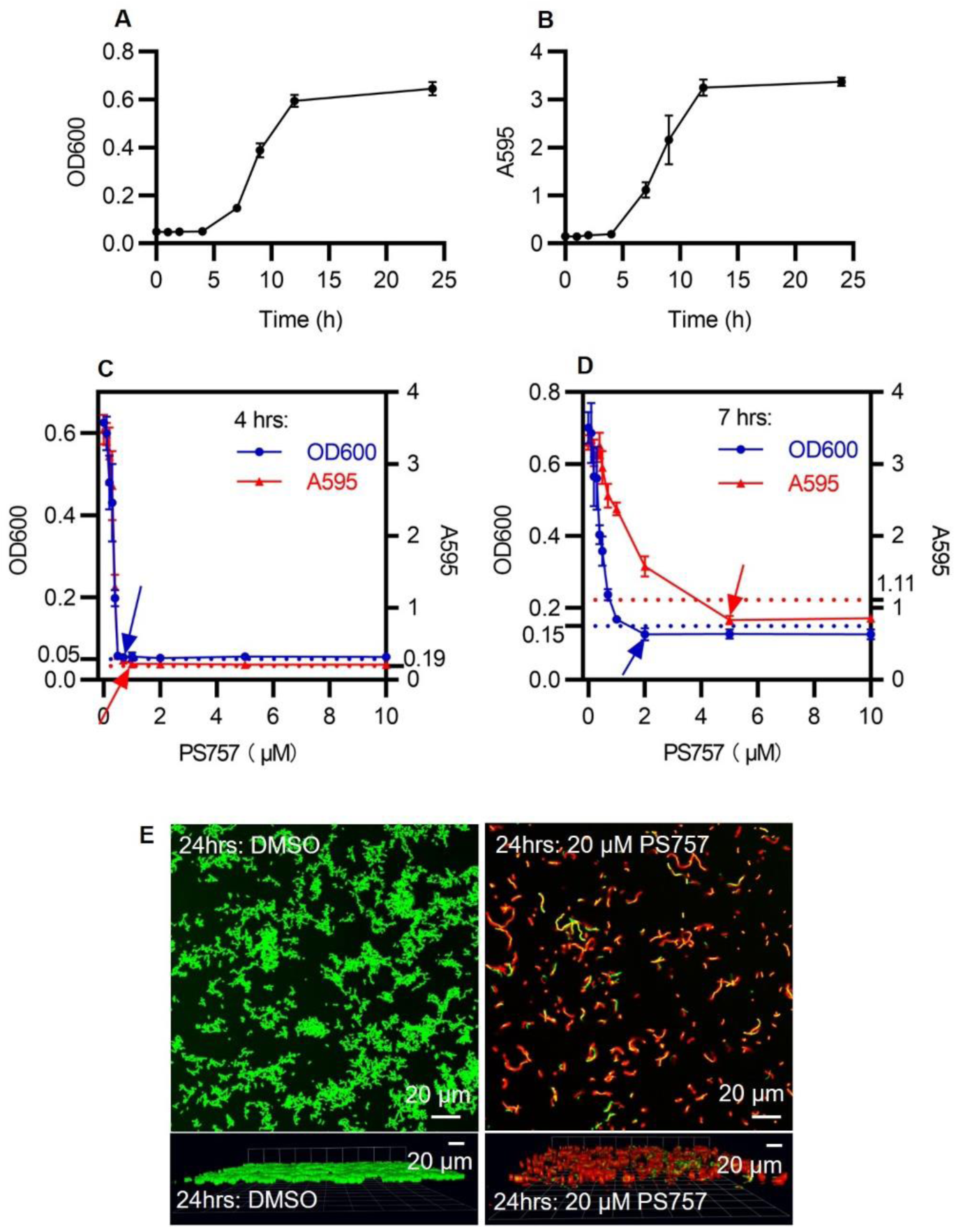
GmPcide PS757 was active against *S. pyogenes* biofilm and caused nucleoid and cell wall abnormalities in *S. pyogenes* cells. (**A & B**) Bacterial growth **(A)** and biofilm formation **(B)** of *S. pyogenes* HSC5 strain were measured in microplate assays for 24 hrs using BHI medium, which identified 4 hrs, 7 hrs, and 24 hrs as the time points for three different phases of *S. pyogenes* biofilm formation, including biofilm initiation, biofilm development, and fully mature biofilm. **(C)** At 4 hrs, PS757 treatment at the concentrations of 0.7 µM and 1.0 µM prevented *S. pyogenes* HSC5 bacterial growth and biofilm formation in the initiation phase, respectively. **(D)** At 7 hrs, PS757 treatment at the concentrations of 2.0 µM and 5.0 µM ceased *S. pyogenes* HSC5 bacterial growth and biofilm formation in the maturing phase, respectively. **(E)** At 24 hrs, PS757 treatment at the bactericidal concentration of 20 µM eradicated mature *S. pyogenes* HSC5 biofilm.

**Fig. 3.**
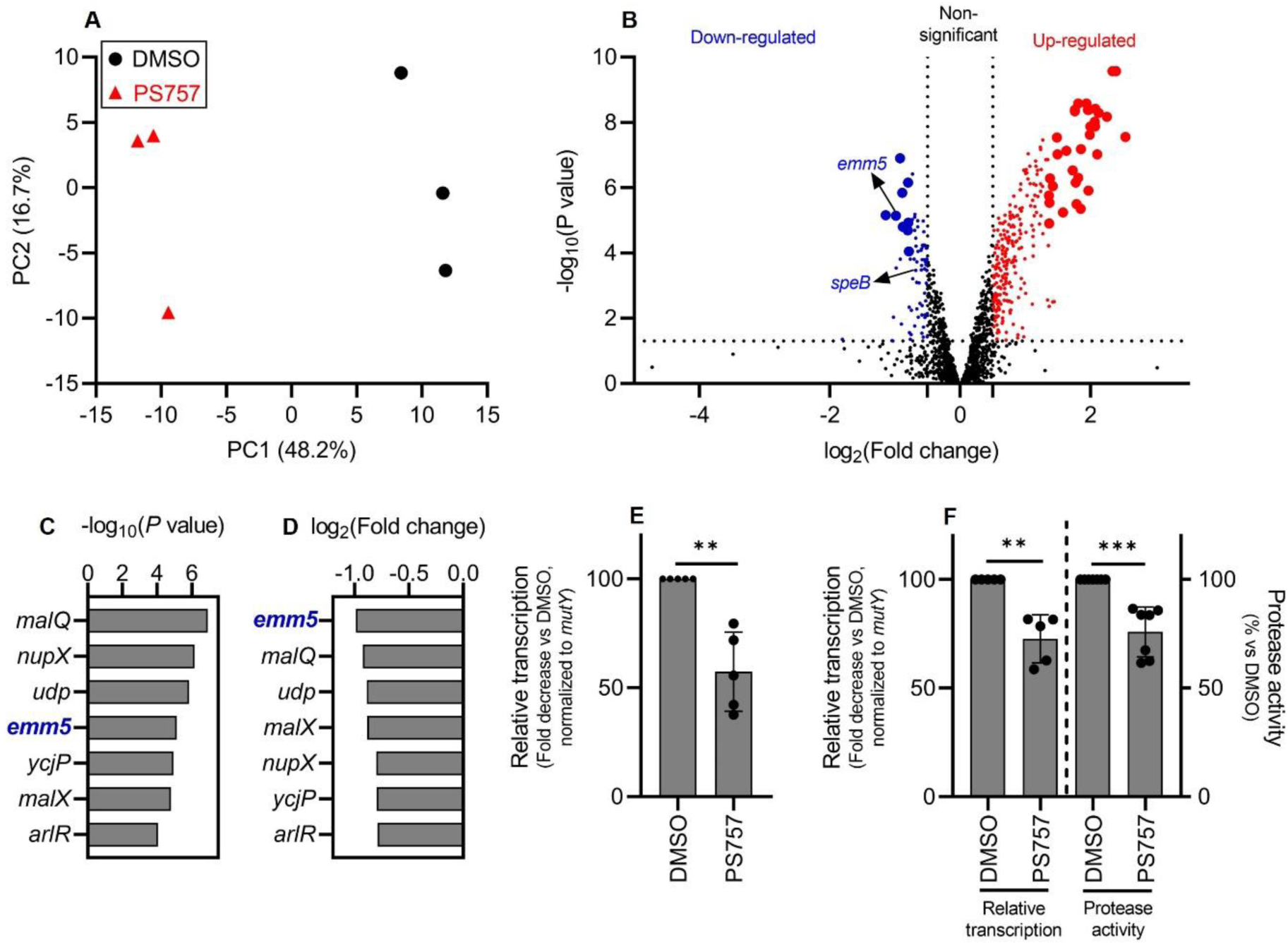
GmPcide PS757 treatment of *S. pyogenes* resulted in altered transcriptome featuring the inhibition of two major virulence factors (*emm5* and *speB*). (**A**) Principal component analysis (PCA) was conducted for comparing the transcriptomes of *S. pyogenes* HSC5 (obtained from RNA-Seq) between two conditions, under the treatments of 0.4 µM PS757 or DMSO vehicle. Score plot (PC2∼PC1) of the first (PC1) and second (PC2) principal components revelaed the transcriptome differences between these two conditions, with clear separation observed along the PC1 axis (accounting for 48.2% of total variance between specimens). **(B)** A volcano plot comparing the PS757-treated vs vehicle (DMSO)-treated was used for identifying differentially expressed genes (DEGs) in RNA-seq analysis. DEGs with log_2_(FC) > 0.5 and P < 0.05, including down-and up-regulated genes, were indicated as blue and red dots, respectively. A more stringent criteria, upper limits of the 99% confidence intervals (CI) for log_2_(FC) and -log(P) among DEGs, was further applied to identify the most significant DEGs, with down- and up-regulated genes indicated as bigger blue and red dots, respectively. **(C & D & E)** Seven genes were identified as the most down-regulated group of genes by the two more stringent criteria (**C & D**). Among which, gene *emm5*, a major and multifunctional virulence factor of *S. pyogenes*, was identified with the highest inhibition among all genes by PS757 treatment **(D)**, followed by validation by RT-qPCR test **(E)**. **(F)** Another major *S. pyogenes* virulence factor, *speB*, was also identified with down-regulated transcription induced by PS757 treatment, which was validated by RT-qPCR and protease activity assays. Statistics were performed with Mann-Whitney U test. P ≤ 0.05 is considered as statistically significant. *P ≤ 0.05, **P < 0.01, ***P < 0.001, ****P < 0.0001, ns indicates not significant.

### GmPcide PS757 Has Anti-Virulence Properties

A comparative transcriptomic analysis was performed to determine if exposure to a sublethal concentration of PS757 (0.4 µM) alters expression of *S. pyogenes* virulence factors. Following 24 hrs of culture in media supplemented with 0.4 µM PS757 or vehicle only (DMSO), RNA was extracted and processed for RNA sequencing (RNA-seq). A principal component analysis (PCA) (*20*) of triplicate samples for each of the PS757 and vehicle only treatment groups showed distinct separation (Fig. 3A), demonstrating PS757 induced distinct transcriptome differences. Analysis of differentially-expressed genes (DEGs, |log_2_(FC)| > 0.5; P < 0.05) identified 75 down-regulated genes (50 genes with an annotated protein function, 40 genes annotated as encoding a “hypothetical protein”) and 277 up-regulated genes (237 genes with an annotated protein function, 40 genes annotated as encoding a “hypothetical protein”) (Fig. 3B). Of the most significantly regulated genes (|log_2_(FC)| and -log(P) > 99% confidence intervals upper limits), 9 genes were down-regulated (7 genes with annotated protein function, 2 genes annotated as encoding a “hypothetical protein”) and 33 genes were up-regulated (32 genes with annotated protein function, 1 gene annotated as encoding a “hypothetical protein”) (Fig. 3B and Table S1). The group of highly upregulated genes featured two ribosomal protein-associated pathways, Rpl and Rps (fig. S4). The significantly down-regulated gene group identified *emm5* (genomic locus: GPBJDOFB_01691) as the most down-regulated gene (Fig. 3, C and D), which encodes the M protein, a cell wall-associated surface protein that plays multiple roles in *S. pyogenes* infection (*12, 21, 22*). Another prominently down-regulated gene was *speB* (genomic locus: GPBJDOFB_01704) (Fig. 3B), which encodes a secreted cysteine protease and multifunctional toxin (*23–27*). Analysis by quantitative reverse transcription PCR (RT-qPCR) confirmed that PS757 significantly inhibited expression of *emm5* and *speB* (Fig. 3, E and F) and significantly reduced protease activity in culture supernatants (Fig. 3F). Together, these results indicate that PS757 has anti-virulence properties on *S. pyogenes* cells exposed to sub-lethal concentrations.

### GmPcide PS757 Ameliorates Tissue Damage in a Murine Model of SSTI

The well-characterized murine subcutaneous ulcer model of *S. pyogenes* SSTI was used to evaluate the ability of GmPcide PS757 to treat an active infection. In the acute phase of this model, subcutaneous injection of 10^7^ CFU of *S. pyogenes* HSC5 into the flanks of 7-week-old SKH1 hairless mice results in a draining ulcer apparent by 24 hrs post-infection (PI) with peak bacterial burdens, measured as recovered CFU, obtained at around Day 3 PI. To assess the efficacy of GmPcide treatment, infected mice received a subcutaneous injection of PS757 (0.03 mg/25 g) or vehicle (DMSO) adjacent to the site of infection at 2, 24, 48, and 70 hrs PI (Fig. 4A). When examined over 3 days, GmPcide-treated mice experienced less infection-related weight loss on Day 1 and gained more weight over the period of observation (Fig. 4B). By Day 3, treated mice also had considerably reduced tissue damage, as shown by a significant decrease in ulcer area as compared to vehicle-treated controls (Fig. 4, C and D), a significant reduction in bacterial burden of approx. 1 log (Fig. 4E) and reduction in serum levels of pro-inflammatory cytokines TNFα (Fig. 4F), IL-6 (Fig. 4G), but not IL-1β (Fig. 4H). Together, these data show that treatment with GmPcide PS757 has efficacy for improving outcomes over the acute phase of SSTI.

**Fig. 4.**
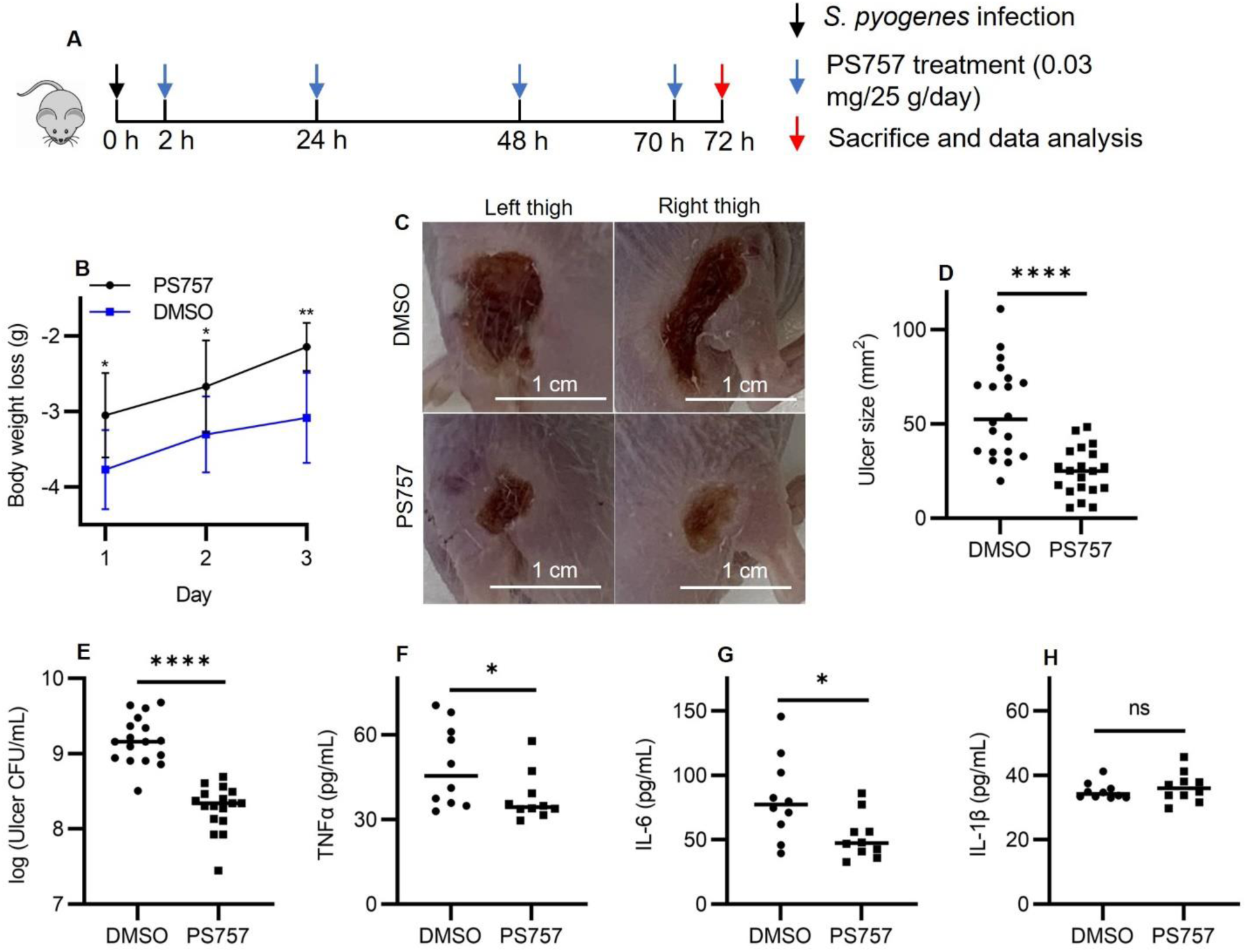
GmPcide PS757 was effective in treating *S. pyogenes*SSTI in mice. **(A)** Timeline of the 3-day infection and treatment protocol by using PS757 to treat *S. pyogenes* SSTI in mice. **(B)** PS757 treatment alleviated acute weight loss caused by *S. pyogenes* SSTI in mice in three days. **(C & D)** PS757 treatment reduced ulcer formation at Day 3 of *S. pyogenes* SSTI in mice (P < 0.0001). **(E)** PS757 treatment attenuated bacterial burden at Day 3 of *S. pyogenes* SSTI in mice (P < 0.0001). **(F & G &H)** Generation of host pro-inflammatory inflammation cytokines, TNFα **(E**, P ≤ 0.05) and IL-6 **(F**, P ≤ 0.05), but not IL-1β **(H)**, were reduced in the PS757-treated group at Day 3 of *S. pyogenes* SSTI in mice. Statistics were performed with Mann-Whitney U test. P ≤ 0.05 is considered as statistically significant. *P ≤ 0.05, **P < 0.01, ***P < 0.001, ****P < 0.0001, ns indicates not significant.

### GmPcide PS757 Treatment Accelerates Healing Kinetics

To test the effect of GmPcides following the acute phase, mice were treated with PS757 or vehicle alone as described above, but were then monitored over a period of 12 days (Fig. 5A). When tissue damage was examined, ulcers in vehicle-treated mice obtained a maximum area at approximately 6 days PI, characterized by the formation of a hard eschar consisting of dry necrotic tissue on the lesion surface (Fig, 5, B and C). In contrast, ulcer area in PS757-treated mice reached maximum area at Day 3 and starting from Day 2 were significantly smaller than those in vehicle-treated mice (Fig. 5, B and C). Furthermore, while the ulcers in the vehicle control group maintained a relatively constant size, those in the PS757 group diminished in size over the course of 12 days (Fig. 5, B and C). Evidence of accelerated healing in PS757-treated mice came from examination of eschars, where in the vehicle control group, only 21% of eschars (4/19 mice) had sloughed off the ulcer surface with new and healthy skin underneath by Day 12 (Fig. 5D). In contrast, eschars began sloughing off from PS757-treated mice starting at Day 6 and by Day 12 75% of mice had lost their eschars (15/20), which were replaced with new and healthy skin (Fig. 5D). PS757 treatment also promoted clearance of the infection with an average 1.3 log reduction in bacterial burden compared to vehicle treated controls (Fig. 5E). These findings demonstrated that PS757 can both limit the degree of tissue damage, as well as accelerate bacterial clearance and wound-healing in *S. pyogenes* murine SSTI.

**Fig. 5.**
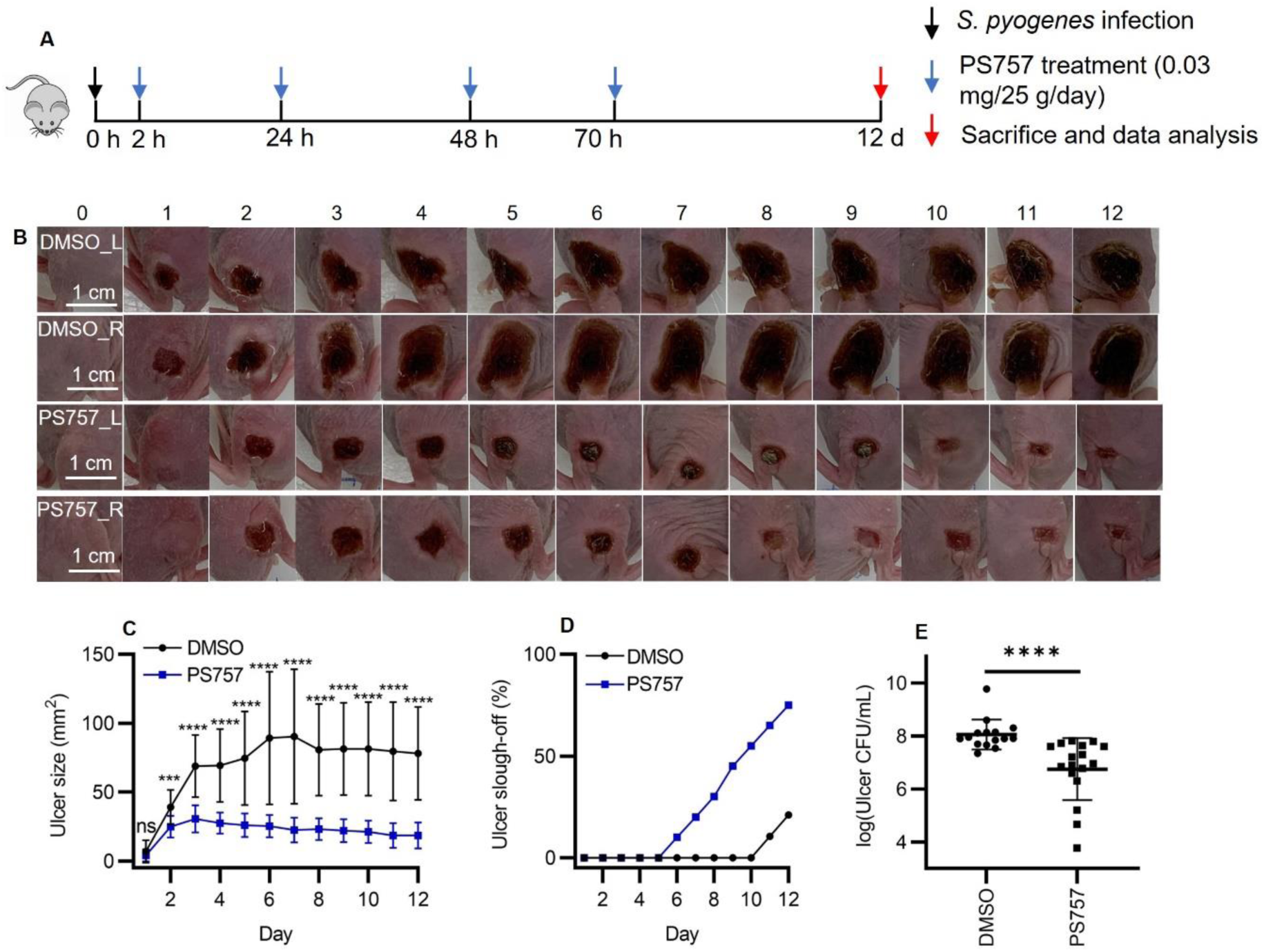
GmPcide PS757 treatment promoted ulcer healing and bacterial clearance in *S. pyogenes* SSTI in mice. **(A)** Timeline of the 12-day infection and treatment protocol by using PS757 to treat *S. pyogenes* SSTI in mice. **(B & C)** PS757 treatment promoted ulcer healing in *S. pyogenes* SSTI in mice during 12 days of infection (P < 0.001). **(D)** PS757-treated mice were observed with quicker eschars slough-off from the infected skin than the untreated group in 12-day S. pyogenes SSTI in mice. **(E)** PS757 treatment promoted the clearance of bacterial infection at Day 12 in *S. pyogenes* SSTI in mice (P < 0.0001). Statistics were performed with Mann-Whitney U test. P ≤ 0.05 is considered as statistically significant. *P ≤ 0.05, **P < 0.01, ***P < 0.001, ****P < 0.0001, ns indicates not significant.

## DISCUSSION

In this study, we found that GmPcide PS757, a member of a novel family of antimicrobial compounds synthesized around a peptidomimetic ring-fused 2-pyridone scaffold, could prevent *S. pyogenes* biofilm formation and effectively treat *S. pyogenes* SSTI in a murine model. Analyses of *S. pyogenes* treated with PS757 revealed anti-virulence properties that involved inhibiting expression of key virulence factors including M protein and the SpeB protease, and the induction of nucleoid and cell wall damage. PS757’s anti-virulence and anti-biofilm activities likely contributed to its efficacy for treatment of *S. pyogenes* SSTI.

In addition to their anti-virulence activity, GmPcides are bacteriostatic and/or bactericidal against a broad range of Gram-positive species (*5*), however, their mechanism of action is unknown. Characterization of activity against *Enterococcus faecalis* has shown that GmPcide are bacteriostatic against exponentially growing cells, but bactericidal against stationary phase cells (*5*). This latter mechanism resembles the process of fratricide, where activation of the GelE protease and the ATN autolysin leads to the lysis of a sub-population of cells to release DNA for the promotion of biofilm formation (*28*). In the present study, we found that GmPcides were bactericidal against both exponential and stationary phases *S. pyogenes*, revealing its robust bactericidal activity against all phases of streptococcal cells. Moreover, when treated with either sublethal or bactericidal concentrations of PS757, *S. pyogenes* cells were observed with nucleoid abnormalities featuring a condensed and filamentous structure, similar to what has been reported for the fluoroquinolone antibiotic nalidixic acid and the lantibiotic nisin, which can cause DNA condensation and fragmentation by targeting the DNA gyrase in *S. aureus* (*29–31*). Exposure of *S. pyogenes* to a bactericidal concentration of PS757 induced cell wall abnormalities with numerous dense globular structures formed at the periphery of the cell wall, which resembles the blisters and bubbles that appear in the cell envelopes of *Escherichia coli* cells that have been treated with the antimicrobial peptide Gramicidin S (*32*). Thus, similar to these other antibiotics, PS757 antimicrobial activity may be associated with adverse membrane stress caused by inhibition of an essential pathway, including DNA replication (*33*), cell wall biosynthesis (*32, 34*), or production of specific proteins (*35*).

GmPcides are one of only a few classes of antibiotics that can kill non-dividing bacterial cells (*5*). Here we show that they also are one of the few antibiotics that are effective at killing cells in biofilm. It has been shown that up to 90% of *S. pyogenes* strains collected from both invasive and noninvasive infections were able to form biofilm (*36*), that *S. pyogenes* biofilm develops on a variety of host surfaces and tissues, and that cells residing the biofilm matrix exhibit higher antibiotic resistance (*37*). Biofilm-associated *S. pyogenes* infections often result in persistent host carriage and recurrent infections and are a major reason leading to treatment failure following therapy with available standard-of-care antibiotics (*15, 36, 38*). Numerous virulence factors, including the SpeB cysteine protease, are upregulated in biofilm, suggesting biofilm potentially enhances pathogenesis (*14, 39*). Thus, the ability of PS757 to effectively inhibit biofilm development may have played a role in its ability to treat *S. pyogenes* SSTI.

The ability of PS757 to inhibit expression of the SpeB toxin and expression of other virulence factors may also have contributed to its efficacy in treatment of SSTI. Toxins are thought to play an important role in the tissue damage that accompanies SSTI and this is the basis for therapies that combine a bactericidal beta-lactam antibiotic with clindamycin or the newly discovered 2S-alkyne, which have been shown to inhibit toxin expression *in vitro* (*23, 40–44*). The mechanism of clindamycin inhibition of toxin expression may involve its ability to alter expression of several regulators of toxin transcription (*45*). Whether GmPcides inhibit toxin expression by targeting transcription regulators remains to be determined. However, this therapeutic approach is being rendered less effective by rising rates of clindamycin resistance. Another approach has been to use passive immunotherapy with intravenous pooled human immunoglobulin (IVIG) (*44*) which may both neutralize toxins and promote neutralization of antiphagocytic and biofilm-promoting virulence factors such as M protein. Thus, the ability of GmPcides to inhibit expression of SpeB, M protein and other virulence factors may have contributed to PS757’s ability to treat SSTI by reducing tissue damage, accelerating bacterial clearance, stimulating ulcer healing, and alleviating host inflammation to promote a quicker recovery. Overall, these results demonstrate that GmPcides hold great promise in preventing and treating *S. pyogenes* SSTI. Our findings will help direct the continuing development and optimization of GmPcide compounds towards a novel class of antibiotics.

## MATERIALS AND METHODS

### Ethics Statement

All animal experimentation in this study was conducted following the National Institutes of Health guidelines for housing and care of laboratory animals and performed in accordance with institutional regulations after pertinent review and approval by the Animal Studies Committee at Washington University School of Medicine (Protocol # 22-0307).

### Bacterial Strains, Culture and GmPcide

All experiments described utilized *S. pyogenes* HSC5 strain (*39, 46*) whose virulence properties in the murine SSTI has been extensively characterized (*8, 9, 47*). Unless otherwise specified, liquid cultures utilized C medium (*39*) and were inoculated from several colonies picked from a C medium plate that had been incubated overnight at 37°C under the anaerobic conditions produced by a commercial anaerobic generator (Becton Dickinson, 260683). Routine liquid culture was performed using 15 ml conical tubes containing 10 mls of media that were incubated under static conditions at 37°C. GmPcide PS757 was synthesized from a 2-pyridone scaffold with as previously reported (*5*) and was chosen for further studies in the current work.

### Determination of MIC and MBC

The minimum inhibitory concentration (MIC) and minimum bactericidal concentration (MBC) represent the lowest concentrations of GmPcide that inhibited growth and viability, respectively. Broth microdilution MIC and MBC assays were performed in BHI medium following the methods established by the Clinical and Laboratory Standards Institute (*5, 48*).

### Determination of GmPcide bactericidal activity against exponential and stationary *S. pyogenes*

Cells from an overnight culture grown described above were harvested by centrifugation, washed with an equal volume of sterile saline (Millipore Sigma, S8776) and resuspended to equal volume of fresh sterile saline. This suspension was used to inoculate fresh C-medium at a dilution of 1:1000, that was then dispensed at 200 μL aliquots into the wells of a 96-well plate (*39*). At 7 hrs and 14hrs post inoculation, exponential and stationary *S. pyogenes* cells were treated with vehicle alone (DMSO) (Sigma-Aldrich, D2650) or with PS757 (20 μM) for an additional 12 hrs of incubation, and CFUs enumerated by quantitative plating.

### Determination of Sublethal GmPcide Concentrations

Cells from an overnight culture grown described above were harvested by centrifugation, washed with an equal volume of sterile saline (Millipore Sigma, S8776) and resuspended to equal volume of fresh sterile saline. This suspension was used to inoculate fresh C-medium at a dilution of 1:1000, that was then dispensed at 200 μL aliquots into the wells of a 96-well plate to which GmPcide PS757 was added at concentrations ranging from 0 to 5.0 µM. The OD600 of the culture was measured using a spectrophometer (Beckman Coulter, DU-800) over the course of 24 hrs, and CFUs enumerated by quantitative plating. The highest functional sublethal GmPcide concentration was determined as the highest concentration of one antimicrobial agent that does not inhibit the streptococcal cells growth.

### Electron Microscopy

Using the microplate assay described above, streptococci were cultured with a sublethal (0.4 µM) concentration of PS757 for 24 hrs or were exposed to a bactericidal concentration by first culturing cells in the absence of GmPcide overnight, followed by the addition of PS757 (20 µM) and continuing the incubation for an additional 5 hrs. Cells were prepared and examined using transmission electron microscopy as described in detail (*49–52*). Briefly, cells were harvested by centrifugation and fixed by the addition 2% paraformaldehyde/2.5% glutaraldehyde (Polysciences Inc.) in 100 mM sodium cacodylate buffer (pH 7.2) to the bacterial pellet for 1 hr at room temperature. Samples were washed in sodium cacodylate buffer and postfixed in 1% osmium tetroxide (Polysciences Inc.) for 1 hr, then rinsed in deionized water prior to en bloc staining with 1% aqueous uranyl acetate (Ted Pella Inc.) for 1 hr. Following several deionized water rinses, samples were dehydrated in a graded series of ethanol and embedded in Eponate 12 resin (Ted Pella Inc.). Sections of 95 nm were cut with a Leica Ultracut UCT ultramicrotome (Leica Microsystems Inc.), stained with uranyl acetate and lead citrate, and visualized on a JEOL 1200 EX transmission electron microscope (JEOL USA Inc.) equipped with an AMT 8 megapixel digital camera and AMT Image Capture Engine V602 software (Advanced Microscopy Techniques).

### Biofilm Culture and Quantitation

Culture in 96-well plates in BHI medium (Thermo Fisher BD, DF0037-07-0) at 37°C were monitored for planktonic growth and biofilm formation by OD_600_ and staining with crystal violet (CV) (*49, 50, 53*), respectively. For the latter, planktonic culture was removed by aspiration and the wells stained with a 0.5% CV solution for 10 minutes, washed with deionized water, air-dried on absorbent paper overnight, and extracted with 33% acetic acid (Sigma-Aldrich, 695092) for 10 minutes.

Extracts were then diluted 20-fold (in 33% acetic acid) and their absorbance at A595 determined using a spectrophotometer (Beckman Coulter, DU-800).

### GmPcide Treatment of Biofilm

Characterization of biofilm cultures indicated that *S. pyogenes* HSC5 biofilm in BHI medium had three distinct phases: initiation, development and maturation; which occurred at 4, 7, and 24 hrs post-inoculation, respectively (Fig. 2B). GmPcide anti-biofilm activity was then determined by the addition of PS757 as follows (fig. S3, A, B, C): At 4 and 7 hrs, PS757 was added to 96-well plate cultures at concentrations ranging from 0-20 μM. At 12 hrs post-inoculation, planktonic and biofilm growth was measured, as described above. For mature biofilm, cultures were prepared using 5 ml of medium in a 35 mm diameter culture dish (MatTek, P35G-0-14-C). At 24 hrs post-inoculation, cultures were treated with vehicle alone (DMSO) (Sigma-Aldrich, D2650) or with PS757 (20 μM) for an additional 5 hrs of incubation. Bacterial viability was then assessed by confocal microscopy using a live/dead fluorescent probe (Thermo Fisher, L7012). Following 30 mins of staining, plates were washed with saline and images acquired using a Zeiss LSM 880 Confocal Laser Scanning Microscope, as previously described (*54–56*).

### RNA Sequencing

Microplate (96-well) culture in C medium was conducted as described above with the addition of 0.4 µM PS757 or vehicle (DMSO). At 24 hrs, multiple wells were harvested and pooled for further processing, with the experiment repeated in triplicate. Extraction of RNA utilized the Direct-zol RNA Miniprep Plus Kit (Zymo Research, R2072) with the quality of the purified RNA determined by spectroscopy (NanoDrop 2000, Thermo Fisher). Libraries for Illumina sequencing were prepared using the FastSelect RNA kit (Qiagen, 334222), according to the manufacture’s protocol and sequences determined using an Illumina NovaSeq 6000. Basecalls and demultiplexing were performed with Illumina’s bcl2fastq software and a custom python demultiplexing program with a maximum of one mismatch in the indexing read. RNA-seq reads were then aligned to the Ensembl release 101 primary assembly with STAR version 2.7.9a (*57*). Gene counts were derived from the number of uniquely aligned unambiguous reads by Subread:featureCount version 2.0.3 (*58*). Isoform expression of known Ensembl transcripts were quantified with Salmon version 1.5.2 (*59*) and assessed for the total number of aligned reads, total number of uniquely aligned reads, and features detected. The ribosomal fraction, known junction saturation, and read distribution over known gene models were quantified with RSeQC version 4.0 (*60*).

### Comparative Transcriptomic Analysis

All gene counts obtained from RNA-seq were then imported into the R/Bioconductor package EdgeR (*61*)and TMM normalization size factors calculated to adjust for differences in library size. Ribosomal genes and genes not expressed in the smallest group size minus one sample greater than one count-per-million were excluded from further analysis. The TMM size factors and the matrix of counts were then imported into the R/Bioconductor package Limma (*62*). Weighted likelihoods based on the observed mean-variance relationship of every gene and sample were calculated for all samples and the count matrix transformed to moderated log_2_-counts-per-million with Limma’s voomWithQualityWeights (*63*). The performance of all genes was assessed with plots of the residual standard deviation of every gene to their average log-count with a robustly fitted trend line of the residuals. Differential expression analysis was then performed to analyze for differences between conditions with results filtered for only those genes with Benjamini-Hochberg false-discovery rate adjusted p-values less than or equal to 0.05. A principal component analysis (PCA) was performed on differential expression data to distinguish differences between conditions (*49*). To find the significantly regulated genes, the Limma voomWithQualityWeights transformed log_2_-counts-per-million expression data was then analyzed via weighted gene correlation network analysis with the R/Bioconductor package WGCNA (*64*). Briefly, all genes were correlated across each other by Pearson correlations and clustered by expression similarity into unsigned modules using a power threshold empirically determined from the data. An eigengene was then created for each de novo cluster and its expression profile was then correlated across all coefficients of the model matrix. Because these clusters of genes were created by expression profile rather than known functional similarity, the clustered modules were given the names of random colors where grey is the only module that has any pre-existing definition of containing genes that do not cluster well with others. The information for all clustered genes for each module were then combined with their respective statistical significance results from Limma to determine whether or not those features were also found to be significantly differentially expressed.

### Quantitative Reverse Transcription Polymerase Chain Reaction (RT-qPCR)

RNA was prepared as described above, with five replicates collected for each treatment condition. Reverse transcription utilized iScript Reverse Transcription Supermix (Bio-Rad, 1708840) with the thermocyler (Applied Biosystems, A24812) programmed as follows: i) Priming for 5 min at 25°C; ii) Reverse transcription (RT) for 20 min at 25°C, and iii) RT inactivation for 1 min at 95°C to acquire complementary DNA (cDNA). After reverse transcription, 12.5 ng cDNA was mixed with primers specific to each gene (Table 2), and PCR conducted using the iTaq Universal SYBR Green Supermix as recommended by the manufacturer (Bio-Rad, 1725121). All qPCR assays were performed on a CFX96 Real-Time System (Bio-Rad), using the following protocol, a 5-minute polymerase activation and DNA denaturation at 95°C, another 10-second DNA denaturation at 95°C, 30 cycles of a 30-second annealing at 60°C, ending with a melt curve with 5 seconds at 65°C first and 5 seconds each at a 0.5°C increase between 65°C and 95°C, with threshold cycles (Ct) obtained at the end of the reactions. Each sample was run in triplicate with average Ct values calculated. Relative expression compared to control was determined by the ΔΔCt method (*65*).

**Table 2.**
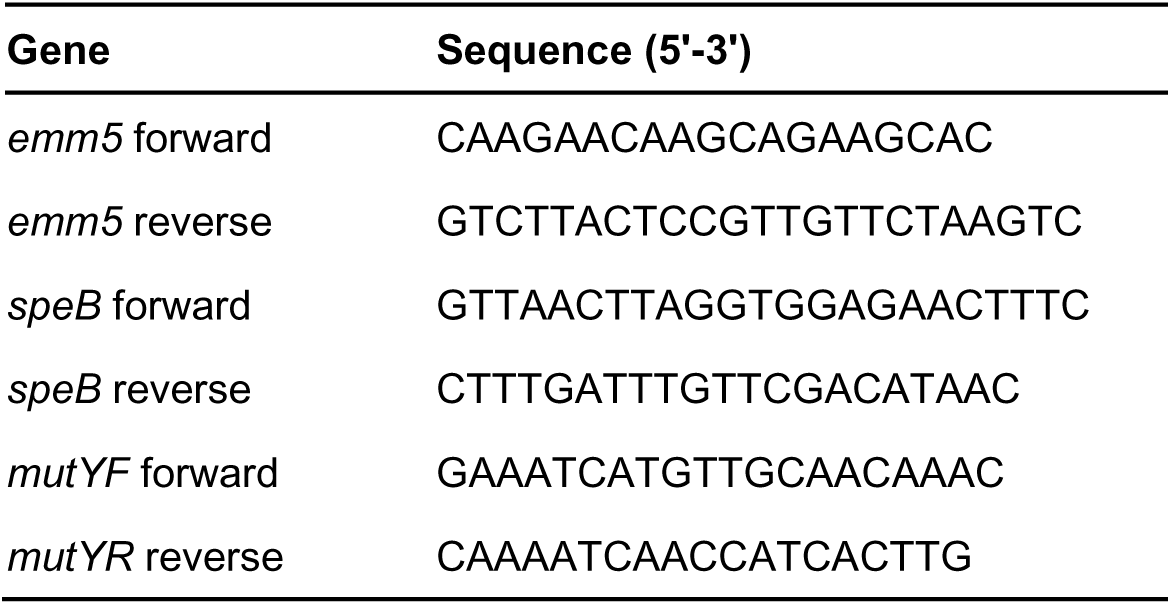
Primers list.

### Protease Assay

Microplate culture was performed as described above in C medium with the addition of 0.4 µM PS757 or vehicle (DMSO). After 24 hrs, extracellular protease activity was determined in bacterial culture supernatant by a method that measures the increase in relative fluorescence generated by the proteolytic cleavage of FITC-casein (Sigma-Aldrich, C0528) (*66–68*). Uninoculated C medium was used to determine background values.

### Subcutaneous Mouse Infection and GmPcide Treatment

Subcutaneous murine infection was conducted as previously described (*8–11*) with the following modifications: For 3-day infections seven-week-old female SKH1 hairless mice (Charles River Labs) were injected subcutaneously with 10^7^ CFU into both the left and right rear thighs, followed by four treatments by subcutaneous injection of vehicle (DMSO) or PS757 (0.03 mg/25 g body weight, dissolved in DMSO) adjacent to the site of infection at 2, 24, 48, and 70 hrs post-infection (PI). Body weight of each mouse was recorded daily and ulcers quantified by digital photography and ImagJ software, as described (*9, 47*). On Day 3, blood was collected using submandibular vein (cheek pouch) method into a collection tube (BD Microtainer, 365967) (*69*). Samples were subjected to centrifugation to obtain serum and levels of selected cytokines quantitated using the DuoSet ELISA kits (*70*), including TNFα (R&D Systems, DY410), IL-6 (R&D Systems, DY406) and IL-1β (R&D Systems, DY401). Mice were sacrificed on Day 3, ulcers were resected and bacterial CFUs determined from tissue homogenates as described (*9*). Initiation of 12-day infection and treatments were conducted as described in the 3-day protocol, no additional treatments were performed and the infection was monitored over the course of 12 days. The data presented are representative of at least two independent experiments, each of which was conducted with 10 mice in each experimental group.

### Data Availability

The RNASeq reads analyzed in this study have been deposited to NCBI data base under the project accession no. PRJNA1040846. The computer codes for the analyses in this study are available in GitHub (https://github.com/QL5001/GASGmPcide_script; branch name, main; commit ID, a27bc81). All data generated or analyzed during this study are included in this published article and supplementary materials.

### Statistics

Unless otherwise stated, conclusions were based on the comparison of means generated from at least three technical and three biological replicates that were tested for significance using Mann Whitney *U* test and conducted using GraphPad Prism 9.0 (GraphPad software). P ≤ 0.05 was considered significant.

## AUTHOR CONTRIBUTIONS

Z.Z., F.A., S.J.H., and M.G.C. conceived and designed the research studies. Z.Z., C.L.P.O., P.S., J.S.P., W.X., and T.M.N. conducted the experiments and acquired data. Z.Z., S.J.H., and M.G.C. analyzed the data. Z.Z. wrote the first draft of the manuscript. Z.Z., K.W.D., S.J.H., and M.G.C. edited the manuscript.

## ACKNOWLEDGMENTS

This work was supported by the NIH RO1DK51406 (S.J.H. and M.G.C.), R01AI134847-01A1 (F.A.), 1U19AI157797-01 (S.J.H., F.A., and M.G.C.), and R21AI163825 (M.G.C.). It was also supported by the Swedish Research Council 2018-04589 and 2021-05040J (F.A.), the Kempe Foundation SMK-1755 (F.A.), and the Erling-Persson Foundation (F.A. and P.S..). Parts of this project were supported under the framework of the JPIAMR–Joint Programming Initiative on Anti-microbial Resistance 2018-00969 (F.A.).

