## Supplementary Materials for "Dihydrothiazolo ring-fused 2-pyridone antimicrobial compounds treat *Streptococcus pyogenes* skin and soft tissue infection"

### SUPPLEMENTARY FIGURES

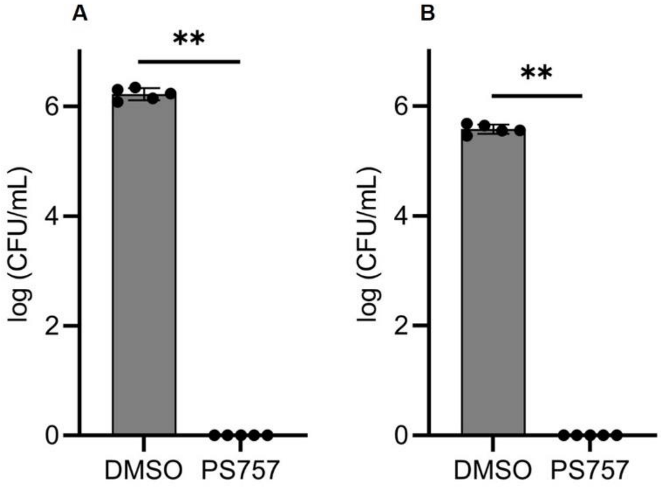


**Fig. S1. GmPcide PS757 demonstrates robust bactericidal activity against the exponential- and stationary-phase of *S. pyogenes* HSC5 cells. (A)** Exponential-phase (7 hours post inoculum) *S. pyogenes* HSC5 cells treated under bactericidal (20 µM) concentration of PS757 for 12 hours were observed with > 6.0 logCFU reduction. **(B)** Stationary-phase (14 hours post inoculum) *S. pyogenes* HSC5 cells treated under bactericidal (20 µM) concentration of PS757 for 12 hours were observed with > 5.0 logCFU reduction. Statistics were performed with Mann-Whitney U test. $P \leq 0.05$ is considered as statistically significant. *$P \leq 0.05$, **$P < 0.01$, ***$P < 0.001$, ****$P < 0.0001$, ns indicates not significant.

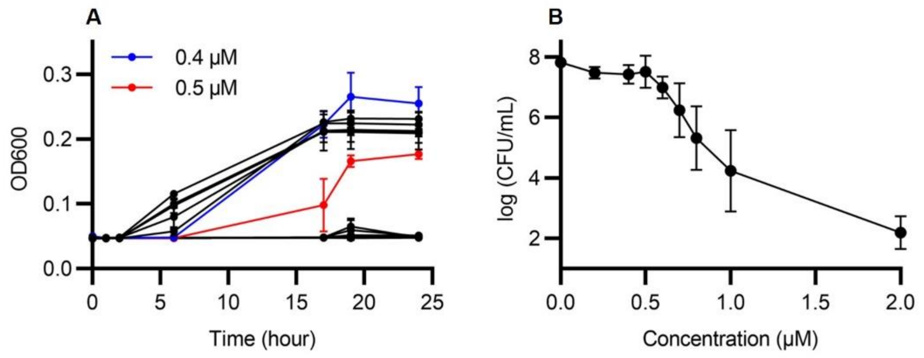


**Fig. S2.** Sublethal concentration of GmPcide PS757 against *S. pyogenes* HSC5 was determined in microplate assay using C medium by measuring both **(A)** OD600 and **(B)** CFU, which identified 0.4 µM as the sublethal concentration of PS757 against *S. pyogenes* HSC5.

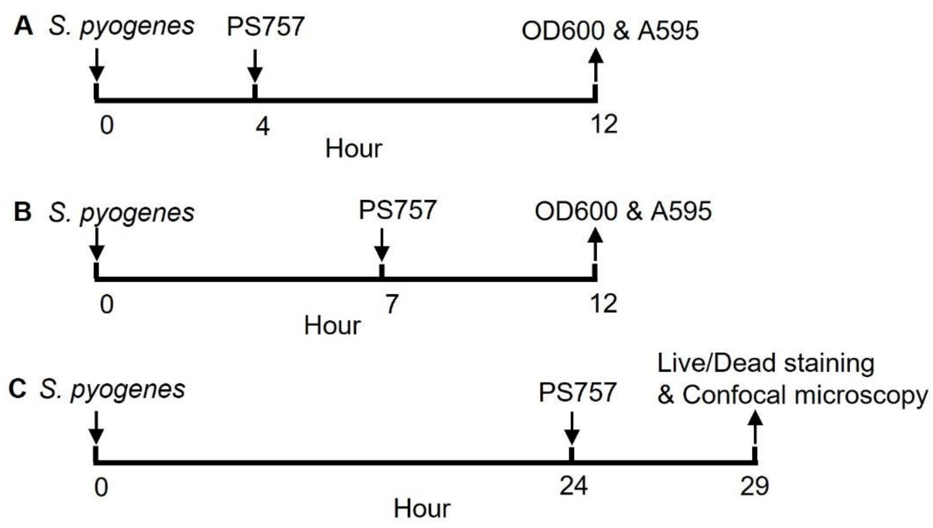


**Fig. S3. GmPcide PS757 treatment to different phases of *S. pyogenes* biofilm. (A)** GmPcide PS757 treatment to *S. pyogenes* HSC5 biofilm at 4 hrs during initiation phase. **(B)** GmPcide PS757 treatment to *S. pyogenes* HSC5 biofilm 7 hrs during maturing development. **(C)** GmPcide PS757 treatment to mature *S. pyogenes* HSC5 biofilm at 24 hrs.

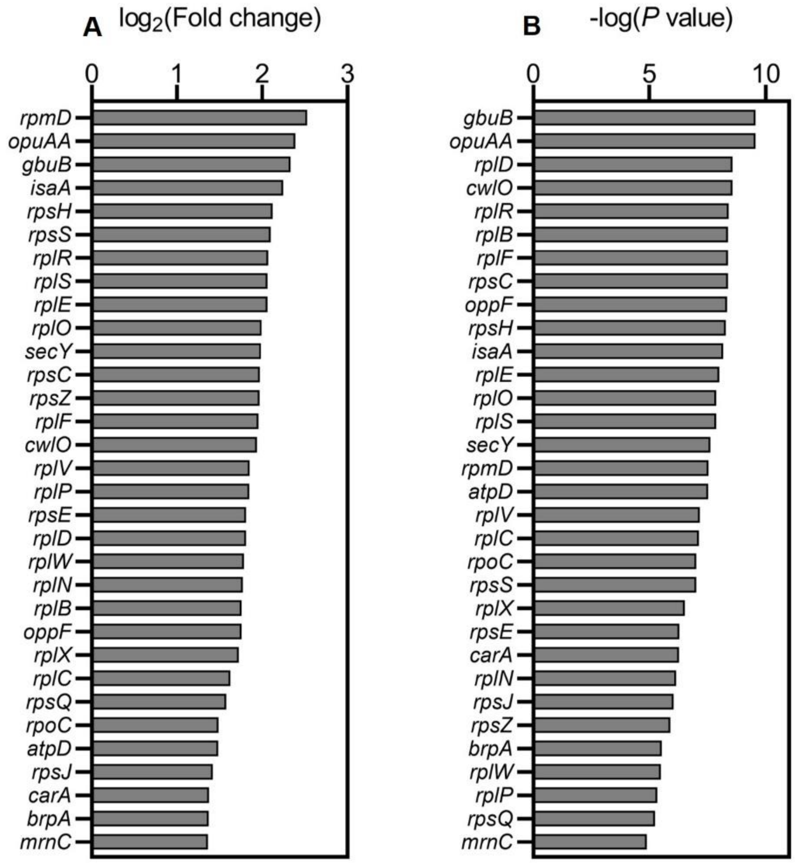


**Fig. S4.** Differentially expressed genes (DEGs) induced by PS757 treatment were identified as $\log_2$(FC) > 0.5 and P < 0.05 in the comparative RNA-seq analysis. Among DEGs, two more stringent selection criteria, |$\log_2$(FC)| **(A)** and -log(P) **(B)** > 99% confidence intervals (CI) upper limits were applied to select the most up-regulated group of genes, which identified 32 most up-regulated genes featuring the involvement of two ribosomal protein-associated pathways, Rpl and Rps.

### SUPPLEMENTARY TABLES

**Table S1.** The list of most down- and up-regulate S. pyogenes genes induced by sublethal PS757 treatment identified by comparative transcriptomic analysis.

| **Differentially expressed levels (DELs)** | **Differentially expressed genes (DEGs)** |
|---|---|
| Most down-regulated | *emm5*, *malQ*, *udp*, *malX*, *nupX*, *ycjP*, *arlR*, *NA*, *NA* |
| Most up-regulated | *rpmD*, *opuAA*, *gbuB*, *isaA*, *rpsH*, *rpsS*, *rplR*, *rplS*, *rplE*, *rplO*, *secY*, *rpsC*, *rpsZ*, *rplF*, *cwlO*, *rplV*, *rplP*, *rpsE*, *rplD*, *rplW*, *rplN*, *rplB*, *oppF*, *rplX*, *rplC*, *rpsQ*, *rpoC*, *atpD*, *rpsJ*, *carA*, *brpA*, *mrnC*, *NA* |
